# Impact assessment and indigenous rights: Cumulative impact assessment of forestry on winter pastures in Mudusjävri reindeer herding co-operative in Finland

**DOI:** 10.1101/2024.10.01.616203

**Authors:** Jan Saijets, Kaisa Raitio, Jarmo Pyykkö, Leena Hansen, Esko Aikio, Pauliina Feodoroff, Osmo Seurujärvi, “What form can an atonement take?” group.

**Affiliations:** Independent Researcher, Tampere, Finland; SLU, Swedish University of Agricultural Sciences, Sweden; Independent Researcher, Inari, Finland; University of Lapland, Rovaniemi, Finland; Independent Researcher, Utsjoki, Finland; Independent Researcher and chairman of Mudusjävri co-operative, Inari, Finland

## Abstract

Assessing the cumulative effects of competing land uses on traditional indigenous livelihoods and culture is a critical component of ensuring the protection of indigenous peoples’ rights. Lack of appropriate methodologies for conducting cumulative impact assessment is a major concern in Sámi indigenous communities across Sápmi, the traditional Sámi territory covering the northern parts of Finland, Sweden, Norway and parts of north-western Russia. We report from a project in which a new type of GIS based impact assessment methodology was developed to assess effects of forestry on winter pastures of the Mudusjävri reindeer herding co-operative in northern Finland. Winter pasture quality was studied as a function of cumulative impacts from forestry. The aim was to develop a way to measure the level of harm caused by forestry on Sámi culture and rights that are protected by national laws and international conventions - in order to assess whether the threshold for significant and hence unacceptable impact has been exceeded. An existing static model for lichen – the main natural winter fodder for reindeer – both for its biomass and growth was used along with mapped forest data to simulate the impacts of forestry on reindeer herding. Our assessment shows that the intensive industrial loggings that started in the 1950s have reduced the ground lichen biomass in Mudusjävri’s pastures by over 30% by 2020. The yearly lichen production has been reduced by 23 - 31% and the yearly lichen depending on the model version used. The respective cost of forestry for Mudusjävri reindeer herding is approximately 370 – 530 000 €/year which is approximately half of the total turnover of the co-operative. Thus, our study indicates that significant harm has been caused on Sámi reindeer herding by other land uses and especially forestry.

## Introduction

Rapidly increasing human induced land use change is an urgent global concern with significant ecological and social impacts [1, 2]. For the world’s indigenous peoples, industrial land use policies pose a threat to traditional livelihoods and the right to practice their culture, as protected by international law [3, 4]. From the indigenous peoples’ rights perspective, one of the critical issues concerns the determination of what constitutes significant, and hence unacceptable, impact [5]. That is, the assessment of whether the impacts of past, present and proposed land uses are about to exceed the threshold beyond which the future of indigenous culture and livelihoods is threatened. In particular, it is extremely difficult for state authorities to carry out their duty to protect and ensure indigenous peoples’ rights to their culture and effective participation in decision-making processes without recognition of the cumulative effects of past, present and planned land uses on indigenous culture, livelihoods and rights. Hence, there is an urgent need to develop assessment practices that capture the cumulative impacts of competing land uses in a way that is relevant and accurate for the livelihoods and aspirations of the concerned indigenous communities and peoples [6, 7].

Indigenous Sámi reindeer herding presents a case in point. Reindeer herding is a traditional semi-nomadic livelihood that forms the cultural and economic basis of Sámi societies in northern Scandinavia, Finland and north-western Russia. Semi-domesticated reindeer graze on natural vegetation over large tracts of land, and reindeer herding is dependent on these natural pastures for its survival. Modern land use forms such as forestry, mining, tourism, hydropower, wind farms and other infrastructure have rapidly expanded to Sámi traditional territories creating detrimental, cumulative effects on the natural pastures of reindeer [8, 9]. Lack of appropriate methods to assess and regulate these effects in land use planning and permitting processes of projects is increasingly highlighted as a threat to Sámi self-determination and the future of Sámi culture [10–12].

In the Finnish part of Sápmi – the traditional territory of the Sámi - the most significant impacts have to date been caused by industrial forestry carried out primarily by the state on so-called public lands. So far, there have been no methods in place to systematically document and assess the cumulative effects of forestry on reindeer pastures when determining the scale and location of forestry operations. This has contributed to persisting and unresolved conflicts between the government-owned forest enterprise Metsähallitus (Finnish Forest Service) and Sámi reindeer herding communities. One of the most central disputes has concerned precisely the severity of the impacts, and whether the impacts caused by forestry activities already executed are so significant that the threshold for significant harm to Sámi culture and livelihoods – as defined in several national regulations - has been exceeded [13, 14].

In this paper we present, to our knowledge, the first ever retroactive cumulative impact assessment of forestry on reindeer winter pastures, the critical bottleneck for reindeer survival. We used a static model for ground lichen biomass [15] developed at Luke, Natural resources institute Finland, and lichen growth dependences [16] formulated by Luke and Helsinki University. The purpose of the paper is twofold: first, to simulate retroactive forestry impacts on reindeer pastures in a way that is accurate and meaningful from a reindeer herding perspective. Second, to present empirical results showing the significance of such impacts in a concrete case of one Sámi reindeer herding community. The study focuses on the change of forest structure in Mudusjävri reindeer herding co-operative’s 2266 km^2^ large area in Inari Municipality in northernmost Finland (Fig 1) and extends from the start of industrial forestry in the 1950s to present time, year 2020.

**Fig 1.**
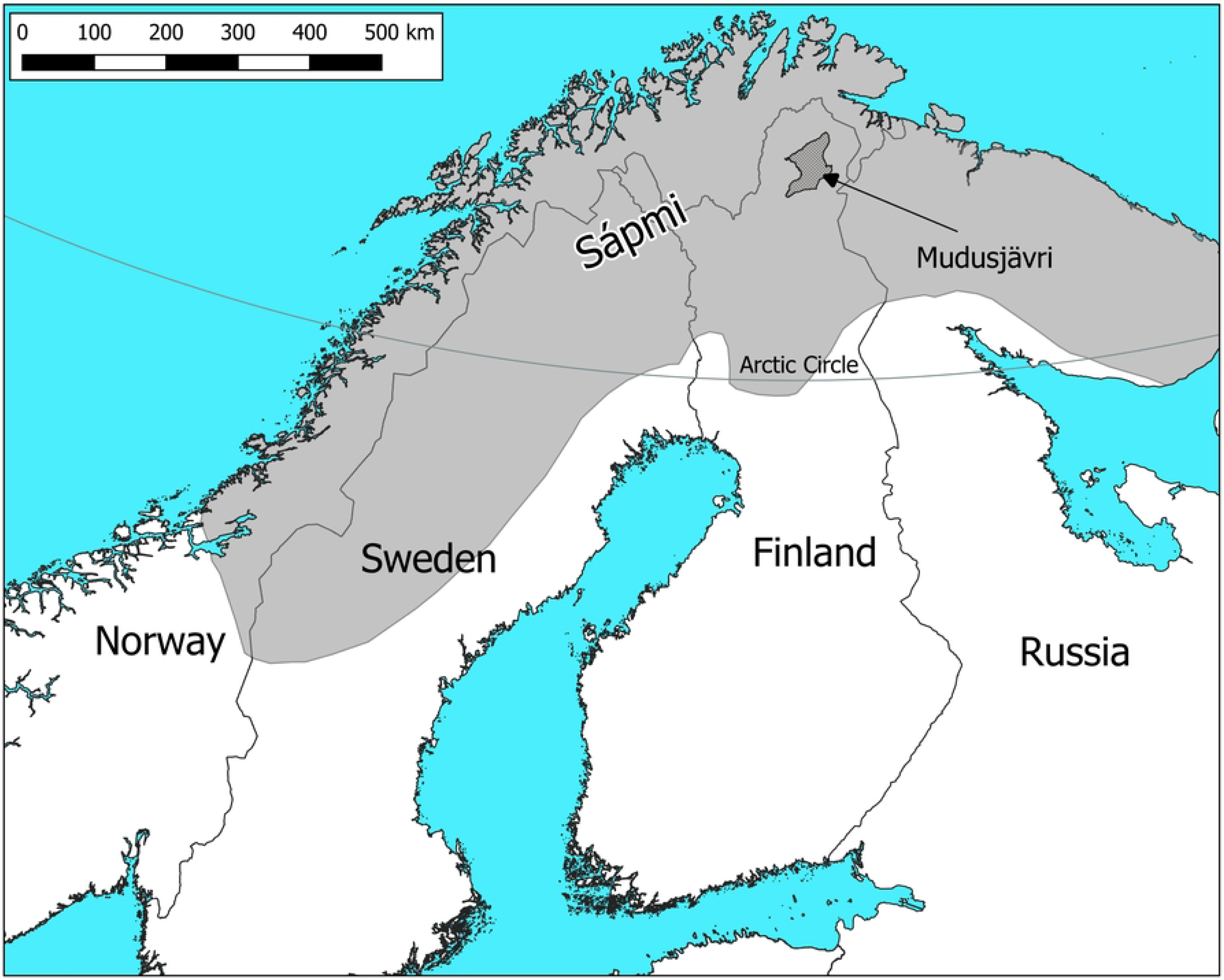
Location of the study area, Mudusjävri reindeer herding co-operative, in central Sápmi.

Ground or terrestrial lichens is the most important mid-winter food for reindeer [17–19] whereas arboreal, tree-hanging lichens are important winter and spring food for reindeer [15, 18, 20].

The pastures were simulated as a function of different parameters with slightly varied models along with the baseline model. The baseline simulation was done by using the lichen biomass and growth models as such as reported in [15] and [16], respectively. The models were also simulated with updated pasture data as well as with a modified model taking into account pasture class change based on Mudusjärvi reindeer herders’ indigenous knowledge on the value of different forest types and individual sites as reindeer pastures. This knowledge was combined with topographic maps, orthorectified aerial infrared photographs, canopy height models, forest resource data, satellite imagery and 52 000 gps-referred photographs into a cartographic method to estimate changes in pasture quality. The assessment of impacts was based on comparing past and present forest structure and its impact on amount and production of lichen, the critical winter food of reindeer. The results demonstrate the significance of the impacts, and highlight the inclusion of herders’ indigenous knowledge as a critical component in developing appropriate cumulative impact assessment procedures. There are dozens of Sámi reindeer herding communities across northern Finland and Sweden being affected by forestry, who can benefit from the presented methodology that captures the impacts of forestry on their livelihood and, consequently, on Sámi culture and rights.

Although, forestry causes several types of negative effects on reindeer herding [21, 22, 23, 24], this study concentrated on the cumulative impacts on lichen productivity. One serious other impact of loggings is the disappearance of arboreal, tree-hanging lichen which is a very important springtime fodder resource for reindeer [15], [18] and can be the bottleneck of economical sustainability of reindeer herding. Other effects are i.a. that reindeer tend to avoid dense sapling stands during wintertime and prefer old forests for grazing as poor visibility in young pine stands might increase predation risk [25], disturbances of tourism, gold digging and infrastructure that tend to drive reindeer to more quiet pastures [26]. These effects combined with the fragmentation of the pastures significantly increases the herding work of reindeer herders as shattered pastures do not keep the herd still.

In the following, we first provide some more background to Sami rights and reindeer herding as a practice. We then present the existing lichen model and the methodology to combine it with our GIS mapping methodology to make lichen simulations. After presenting the results, we discuss the accuracy, validity and limitations of our approach and end with some general conclusions as to the implications of the study in Sápmi and beyond.

### Sami rights, reindeer herding and cumulative effects

The legal status of the Sámi as indigenous people in Finland guarantees the right to practice their culture, including traditional livelihoods such as reindeer herding, hunting and fishing. This right is codified both in the Constitution as well as in several pieces of specific legislation concerning the Sámi and/or natural resources and the environment [27, 28]. Finland is also bound by several international treaties to the same end, most notable being the Covenant on Civil and Political Rights and its paras 1 and 27 (ibid).

However, despite the formal recognition, their rights, reindeer herding and other traditional Sámi uses of land and waters are in many areas heavily affected by cumulative (i.e. combined) effects of competing land uses, e.g. via landscape fragmentation and disturbance of the reindeer [8, 29]. Under such circumstances, it is extremely difficult for state authorities to carry out their duty to protect and ensure indigenous peoples’ rights to their culture without a robust assessment of the cumulative effects of past, present and planned land uses on Sámi culture, livelihoods and rights. This is due to two reasons. First, it is not possible for the affected Sámi reindeer herding communities (*paliskunta* in Finnish, *paalgâs* in Aanaar Sámi language) or other affected Sámi actors to make a decision about a proposed development without knowing the full extent of its impact. In other words, the right of affected Sámi indigenous communities to grant or withhold their Free, Prior and *Informed* Consent (FPIC) as defined in international and national law, is contingent on an adequate and robust assessment of the cumulative impacts on Sámi livelihoods, culture and rights. Second, the responsible public authorities are dependent on an adequate cumulative impact assessment in order to be able to assess whether future projects – such as logging of forests - will cause significant harm to Sámi livelihoods, and hence should not be allowed. This is paramount for ensuring compliance with the Sámi right to culture, as guaranteed in national and international law. Unless the baseline of existing impacts is accurately established, the state risks failing its duties by allowing activities that would together with earlier activities create significant harm.

In fact, UN Human Rights Committee has in 2019 specifically requested Finland to “clarify how the concept of “significant harm” is defined and applied in practice when assessing the impact of measures, including development projects that may directly or indirectly affect the Sámi culture and traditional livelihood”. (CCPR/FIN/QPR/7, April 2^nd^ 2019, para 23). As late as in April 2020, the Committee was, based on Finland’s response in the matter, concerned over the vagueness of the methods used to assess impacts, and urged the state to review existing legislation, policies and practices to ensure a meaningful FPIC process (CCPR/C/FIN/CO/7, April 1^st^, 2020 paras 42 and 43). The purpose of the presented study is to contribute to this end by presenting one method how such assessments can be carried out.

### Methodology and data sets

Our method was to use existing lichen biomass [15], growth [16, 30] and usability [16] models for simulating the lichen productivity of Mudusjävri reindeer herding co-operative’s pastures using accurate GIS-data. The original lichen biomass model was used as the basis which compared to five model variations based newer pasture inventory data and reindeer herders’ traditional knowledge on the pastures.

The dependence of lichen production on forest structure has been established in numerous earlier studies [15, 16, 21, 25, 30–36] and inventories of reindeer pastures nationally [31–34]. This dependence is described in the static lichen biomass model of [15] which is dependent on many parameters like reindeer amounts, forest stand development class, infrastructure disturbances, grazing area type and trophic level of site. The model is rewritten here in a simplified form. The lichen biomass can be calculated by equations

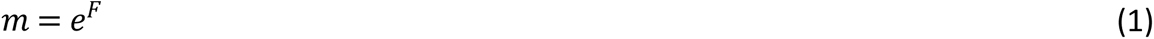

Where *F* is a linear equation

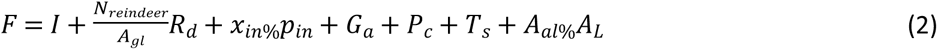

*I* is an empirical constant (*I* value is 6.7) that includes constants with rain and temperature parameters of an average year. *N_reindeer_* is the amount of reindeer winter herd in the co-operative, *A_gl_*, the total area of dry trophic level of sites in square kilometers, *R_d_* value is −0.07 and is a model parameter for reindeer amount, *X_in%_* is the percentage of co-operative area disturbed by infrastructure, *p_in_*is the model parameter for infrastructure disturbances with a value of 0.032. *G_a_* is a grazing area parameter depending of the area being winter grazing area (*G_a_* = 0), summer grazing area (*G_a_* = – 1.08) or an all-year-around grazing area (*G_a_* = −0.82). Maybe the most important parameter in this study is *P_c_* defining pasture class. Its value dependence is shown in Table 1 for the original fitted model [15] along with some newer values based on the latest pasture inventory [34].

**Table 1.** Pasture class parameters used.

| Pasture class, $P_c$ | Original fitted model [15] | Based on latest inventory [34] |
| --- | --- | --- |
| Old pine forest (> 140 years) | -0.042 | 0 |
| Mature pine forest (81 - 140 years) | 0 | -0.24 |
| Young pine forest (31 - 80 years) | -0.384 | -0.30 |
| Logging areas and sapling stands | -0.370 | -0.55 |
| Mountain birch forests | -0.373 | -0.64 |
| Mountain heaths | -0.608 | -0.52 |

Values of pasture class parameter, $P_c$ .

*T_S_* is the trophic level of site. For dry soil type, the value is 0 while for semi-dry soil type the value is −0.4. AL is the arboreal lichen parameter with a value of 0.02. *A_al%_* is the percentage of arboreal lichen forests of land area and *A_L_* an arboreal lichen parameter with the value of 0.02. This lichen biomass model was empirically fitted to the pasture inventory data of 20 northernmost reindeer herding co-operatives [15] where Mudusjävri co-operative is located.

The yearly lichen production (in kg) of a specific stand is dependent on the biomass and pasture type according to equation [16]

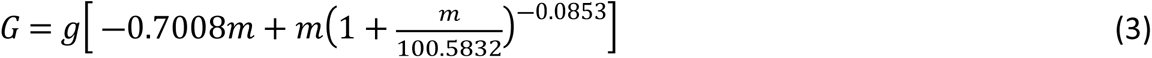

The parameter *g* is the growth factor dependent on pasture class being 1 for and old or mature forest stand, 0.6 for logged or young forests or mountain birch forests and 0.4 for mountain heaths. The fraction of lichen that reindeer can effectively use from a specific site is dependent on biomass of the stand and is described by [16]

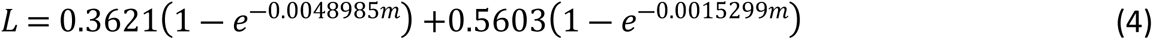

Combining these equations or lichen biomass and productivity models with an accurate pasture GIS-analysis of Mudusjävri co-operative area a cumulative impact assessment on land use change on lichen productivity can be made. This model produces average lichen amounts, growth and usability amounts in pasture classes and do not represent the real amounts of pasture stands. We implemented the pasture simulation as a python code reading a GIS shape-file dbf-table file and calculating lichen biomass, yearly production and usability for each GIS polygon area for the period 1950 to 2020.

To assess the impacts of forestry on reindeer herding based on this relationship, the study consisted of four research tasks

1. Categorize the forest stands in Mudusjävri reindeer herding co-operative into relevant categories based on their value as reindeer pastures
2. Map the history of forestry on each stand from 1950s to 2020
3. In addition to the baseline model, make slightly varied lichen production models based on updated pasture inventory data and reindeer herders’ traditional knowledge
4. Assess the retroactive impacts of forestry on Mudusjävri pasture production based on steps 1-3 by using the simulation model with GIS-data

These steps combined multiple data sets consisting of pasture inventory data; topographic maps, orthorectified aerial infrared photographs; canopy height models; forest resource data; satellite imagery; in situ photos of the forest stands and reindeer herders’ indigenous knowledge both on the value of different forest types as pastures as well as of the characteristics of different stands *in situ*. A summary of the methodology and data used in each step of the process is presented in Table 2. Each step is described in detail in the following sections of this chapter.

**Table 2.** Summary of study methodology.

|  | Research task | Data |
| --- | --- | --- |
| **1** | **Categorizing the forest into relevant stands from a reindeer pasture perspective** |  |
|  | Separating forest polygons from other water and swamps | Topographic maps, National Land Survey of Finland (NLSF) |
|  | Categorizing each forest stand based on their height and density | Orthorectified aerial infrared photos (NLSF)<br>Canopy height model (Forest Centre) |
|  | Classification of low productive forest and land pasture classes | Pasture class data from Luke, Natural Resources Institute, and Syke, Finnish Environment Institute [37] |
|  | Mapping of the disturbance areas and historical pastures and their trophic level of sites based on reindeer herders' traditional knowledge | Reindeer herders' knowledge of the pastures and of reindeer needs |
| **2** | **Mapping the forest history and the progression of forestry** | Historical aerial images from 1960 and 1969 (NLSF)<br>Landsat satellite imagery 1971 – 2016 (NASA) |
|  |  | Sentinel satellite imagery 2016 – 2019 (EU/ESA) |
|  |  | Interviews with a logger employed 1958 – 1972 (to report on forestry prior to 1971) |
|  | Calibration of map data with on-site photographs | 52000 gps-referred images photographed between 2018 and 2020 |
|  | Calibration of map data through reindeer herders' interviews |  |
| **3** | **Making of varied lichen production models based** | Newer pasture inventory data and reindeer herders' traditional knowledge |
| **4** | **Assessing the impacts of forestry on pasture quality** |  |
|  | Using the lichen production model to assess the lichen production in today's forest structure and in the unmanaged forests prior to industrial loggings |  |
|  | Combining pasture statistics with the logging area amounts from 1940 to 2020 to estimate the cumulative impacts of forestry |  |
|  | Using the prices of supplementary fodder to calculate the economic value of the loss in pasture quality and productivity |  |

Summary of the methodology developed in the study, and the data used in each step of the process.

### Categorizing the forest stands into relevant categories based on their value as reindeer pastures

Using the linear lichen biomass and productivity models require the mapping of different forest development classes to be defined as model pasture classes. To be able to digitize or define the forest polygons to be used for area analysis, we used the following background data.

1. Topographic maps, National Land Survey of Finland (NLSF) [38]
2. Orthorectified aerial infrared photos, NLSF [38]
3. Canopy height model, Forest centre [39]

Topographic maps were used as the basis to determine forest areas whereas canopy height models and aerial photos were used for finding logging areas as shown in Fig. 2. This digitizing was best done manually and logging areas were divided into logging areas typical in the co-operative area:

- Logging area without saplings
- Older loggings with dense sapling stand
- Old loggings with saplings in clusters with open spaces between
- Thinned forests stand, tree height over 8 m
- Thinned forest stand, tree height below 8 m
- Old cautious selective logging

**Fig 2.**
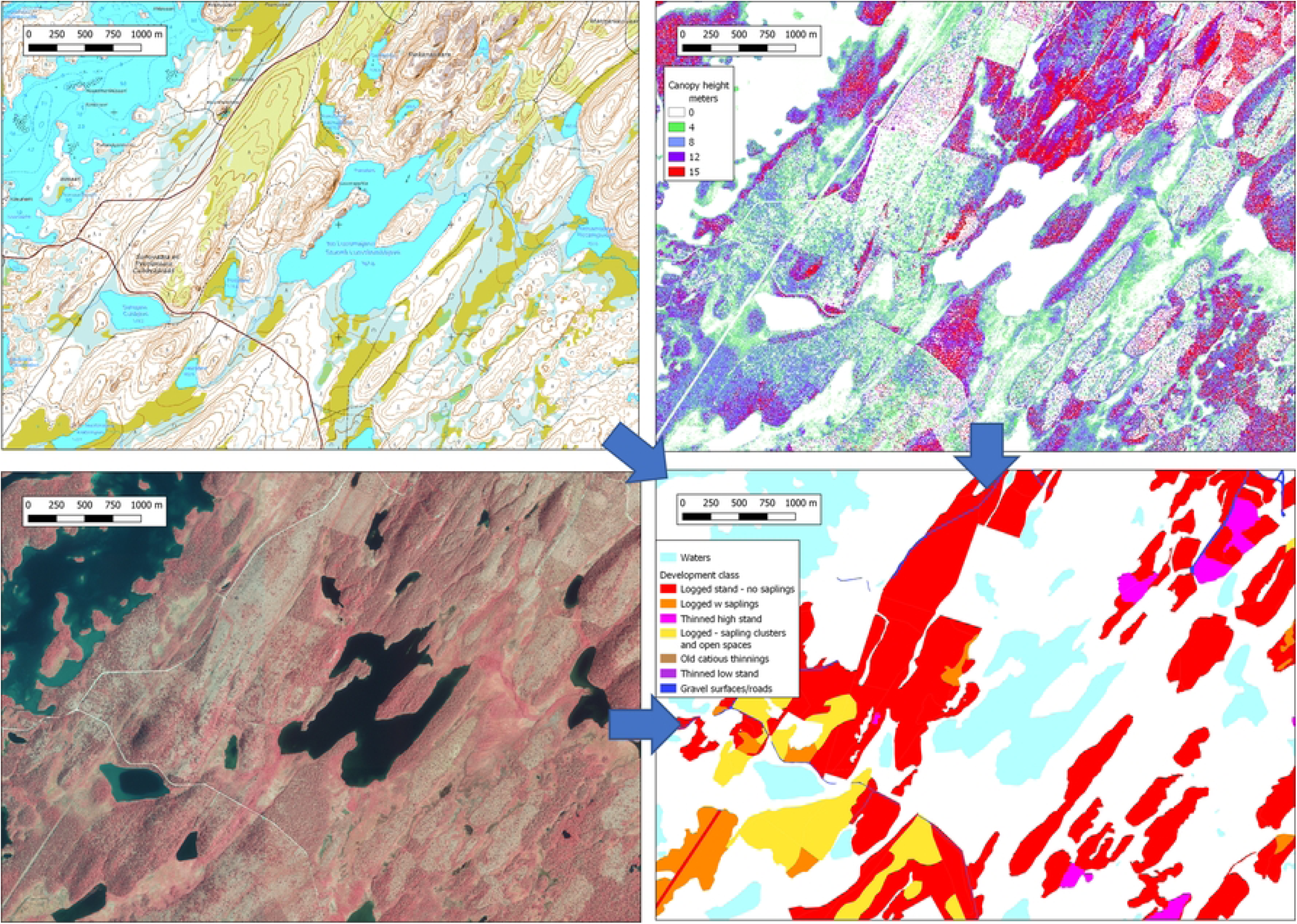
Digitizing logging areas from topographic maps, canopy height models and infrared aerial images.

All of the data and imagery are publicly available and the download sites are referred in the above numbered list. All of the data are typically in a format that can be used by geographical information system (GIS) software.

The topographic maps are available in vector format or in raster formats optimized for different map scales. Orthorectified aerial photos are available both in normal colours or in false-colour infrared images. The orthophoto resolution is 0.5 m/pixel and are enough for our purposes. In our work, we used false-colour infrared photos as they show much clearer the difference between logged areas and otherwise open terrain. Also, the colour difference between deciduous and coniferous forest is very clear.

One of the most important data were the tree canopy height models produced out of laser scanning data. The canopy height model bases on laser scanning by NLSF that has been done from a 2 km altitude sending laser pulses with an average distance of 1.4 m [40]. At ground level the pulse beam width is approximately 50 cm. Several reflections and their respective heights are recorded for each laser pulse. The available canopy height data has a resolution of 1 m x 1 m and can be most easily viewed in by assigning colour ramps for height as in Fig 2 using the QGIS-program [41].

The remaining land areas outside of the defined logging areas were defined as natural state forests based on canopy height and density or as low productive forests, swamps, mountain heaths or other shrub lands. Natural forests were digitized with neighborhood analysis in GRASS GIS software [42] and divided into three categories:

- Dense old pine forest stand, over 50 trees over 13 m per hectare
- Mixed tree height old pine forest stand, 20 – 50 trees over 13 m high per hectare
- Young pine forest (productive forest), less than 20 high trees/hectare and average canopy height more than 7 m in a 21 m diameter circular neighborhood

Only the productive forest lands were digitized and the remaining pasture categories of low productive forests, swamps, shrub lands, mountain birch forests and heaths were taken from the pasture class raster map by Natural Resources Institute Finland and Finnish Environment Institute [37]. In case it was difficult to decide whether the area has been logged in the past or not it was defined as natural stand. This approach was decided not to overestimate the effects of loggings in the final analysis. The same area of Fig 2 now with the natural forests is shown in Fig 3.

**Fig 3.**
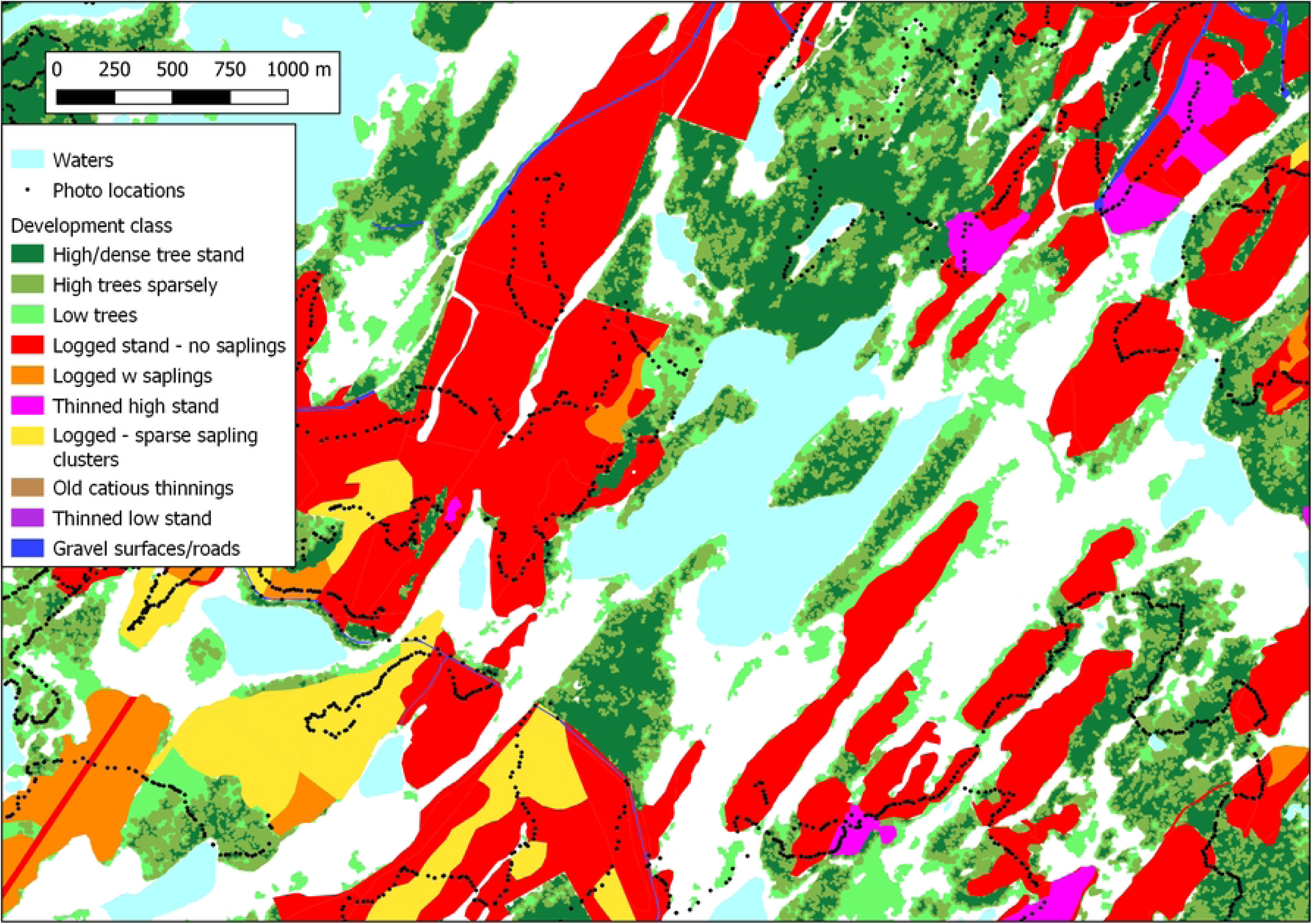
Natural forest polygons determined by canopy neighbor analysis in addition the previously defined logged stands.

### Mapping the forest history from 1950s to 2020

The forestry history was mapped using mainly Landsat satelite imagery (1971 – 2016) from NASA and Sentinel satellite imagery (2016 – 2019) from EU/ESA. These images were downloaded from their respective services [43, 44] in geotiff-format for every spectral band and later combined to proper false-colour infrared images. In QGIS-program this can be done using virtual raster tool.

The resolution of Landsat images are 25 m/pixel and the benefit of using Landsat imagery is the long history of Landsat program. There are images from early 70’s although the resolution of Landsat images was then much lower. Sentinel 2 satellite images can be used in a similar manner as Landsat images but with the benefit of higher resolution of 10 m/pixel. However, the European Sentinel satellites have been up in orbit for only a few years yet and are not useful in tracing earlier forest changes. The oldest Sentinel 2 -satellite images from Mudusjävri region are from 2016 and all earlier logging times were traced from Landsat imagery. Satellite images are not as accurate as laser scan canopy data but rather aid to date the stand logging time as in the example of Fig. 4 that show the forest change between summers 1993 and 1994 just North of Inari village. Rather accurate yearly land changes can be estimated after 1984 using Landsat data as continuous good quality images are available. Earlier images are from 1972 and 1973 after which there is an 11-year gap in history data.

**Fig 4.**
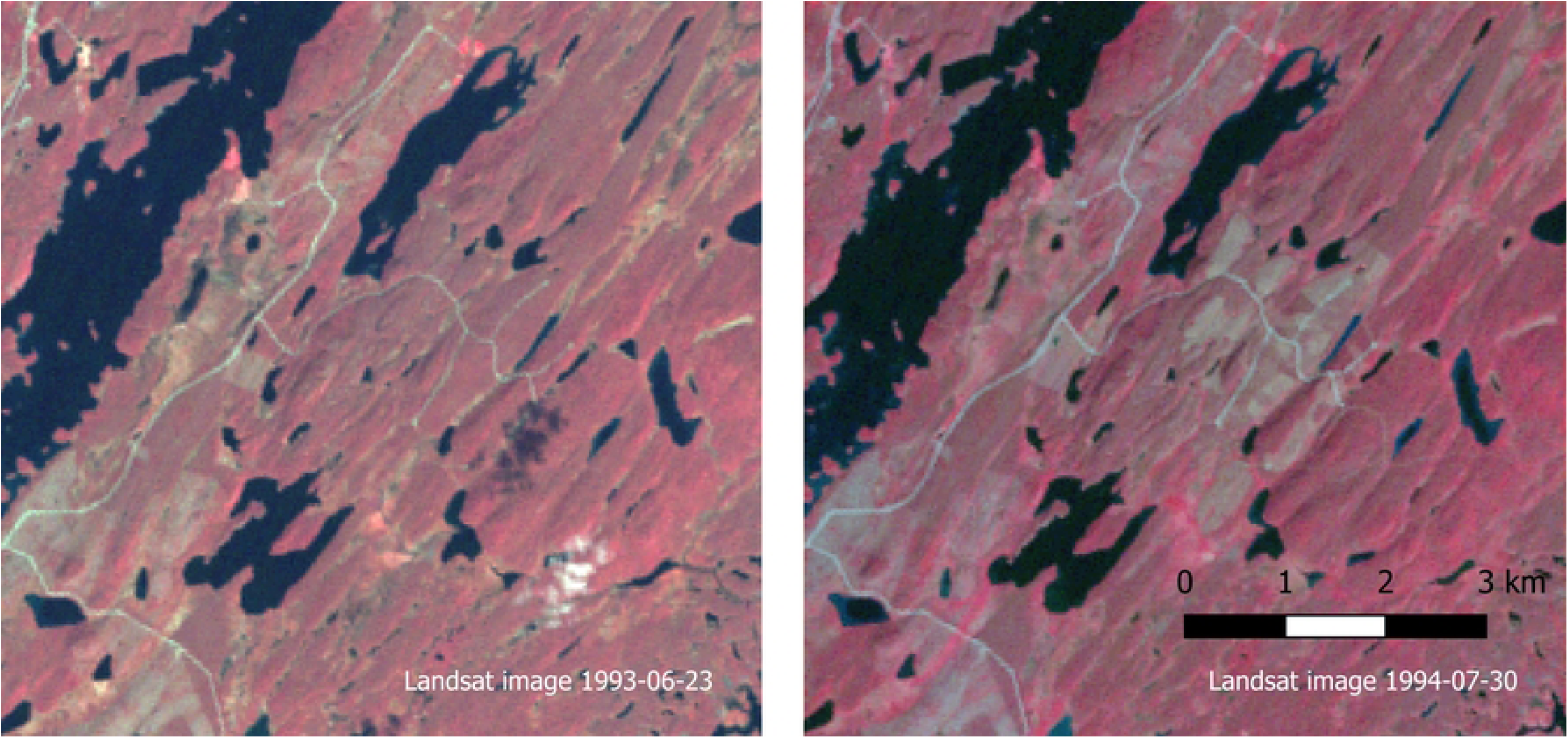
Landsat images of consecutive years are used for tracing forest changes and logging history.

Logging history prior to the 1970s was documented based on NLSF aerial images from 1960 and 1969 and interviews with retired forestry worker who was employed 1958-1972, and are rather rough due to the long time since the events we sought to gain information on. Some 50 % of logging areas prior to 1970s were not possible to time through interviews and the remaining logging areas are seen in 1972 Landsat images were estimated to follow timber felling statistics of the Inari region [45]. Similarly, the 11-year gap between 1973 to 1984 was interpolated following the timber felling statistics. The value of mapping the logging history is that it assures map accuracy and enables modelling of ground lichen amount and growth modelling of the historical pastures.

### Role of reindeer herders’ indigenous knowledge

Close collaboration and knowledge co-production with the herders in the Mudusjävri herding community was essential during several stages of the assessment. First, the categorization of different forest stand types (vegetation types and stages of succession) needed to be based on their value as reindeer pastures. Accurate categorization was dependent on the herders’ knowledge of reindeer preferences and behavior in their area. This categorization needed to be created from the start, because the existing state forest agency categorizations forests had been created to reflect a timber production rationality, and resulted in maps and statistics that were not helpful for the task in hand. Further, the state agency had declined to release any of its data to Mudusjävri herding community and this project.

Second critical role of the herders was to validate the maps created based the digital data, by providing knowledge on the characteristics of different stands *in situ.* Due to their detailed knowledge of the landscape and each forest stand, they were in several cases able to point out inaccurate interpretation of the stand characteristics based on the digital data alone. Old reindeer herders mapped the good quality ground lichen areas before the impacts of forestry. These areas were used for info on trophic level of site described more accurately later in a lichen model variation.

Third, the interviews with herders provided important context for understanding the impacts of forestry in different areas, and in ways that go beyond its impact on pastures to capture the social implications of the pasture change. While this information cannot be captured in maps, we maintain that such qualitative data should be included in any cumulative effects assessment if it is to reflect the impacts of pasture loss and change not only on reindeer, but on reindeer herding as a livelihood and cultural practice (see [46] and [47]). For instance, the impact from the loss of a pasture area depends on its original value as a pasture – some were not high quality to begin with due to their natural characteristics, so logging them would have less impact in comparison to high quality pastures. Further, the severity of the impact depends also on the location of each pasture area within the landscape and thus its function for reindeer herding. Some areas play a more critical role in the seasonal cycle or in collecting the herd and stopping it from spreading into neighboring communities or to large lakes. Due to their functional importance, losing such areas has more severe consequences than one would assume based on their size and productivity alone. Other important areas, on the other hand, may remain, but have lost their value as pasture due to disturbance from e.g. tourism that causes reindeer not to stop in that the area anymore. The interviews showed that due to loss of critical areas – in combination with other factors - some winter groups (*siida*) had seized to exist already decades ago. Such losses to the community and to the continuation of the livelihood in certain areas and families can only be captured through interviews in which the interviewer is able to develop high level of trust in order for herders to disclose such information. We have considered it unnecessary and unethical to disclose details of such knowledge here, but have rather used it to help develop our understanding and to highlight the more general lessons emerging from it, e.g. in terms of differentiated impact based on the function of the different pasture areas.

Finally, our map data of forest characteristics was calibrated and confirmed by taking 52 000 gps-referred photos around critical regions in the co-operative and compared to the produced polygon data. These photos were taken during snowless times in 2018 - 2020 using Pentax K-1 mark 1 SLR for 50 000 photos and Canon 5D mark IV SLR for 2 000 photos. The terrain photo library will be extended during 2020. Care was taken to ensure that GPS tracking was acquired and practical testing was done to measure the location accuracy. Based on those, it seemed that Pentax SLR location accuracy is better than 4 m and Canon SLR is better than 15 m which are both sufficiently good numbers that easily fulfil our required accuracy. An example of previously used map location with gps-tagged photos are shown in Fig 5. Over two hundred workdays were done in the field collecting these GPS-referred photos and 80 % of this was done by one worker.

**Fig 5.**
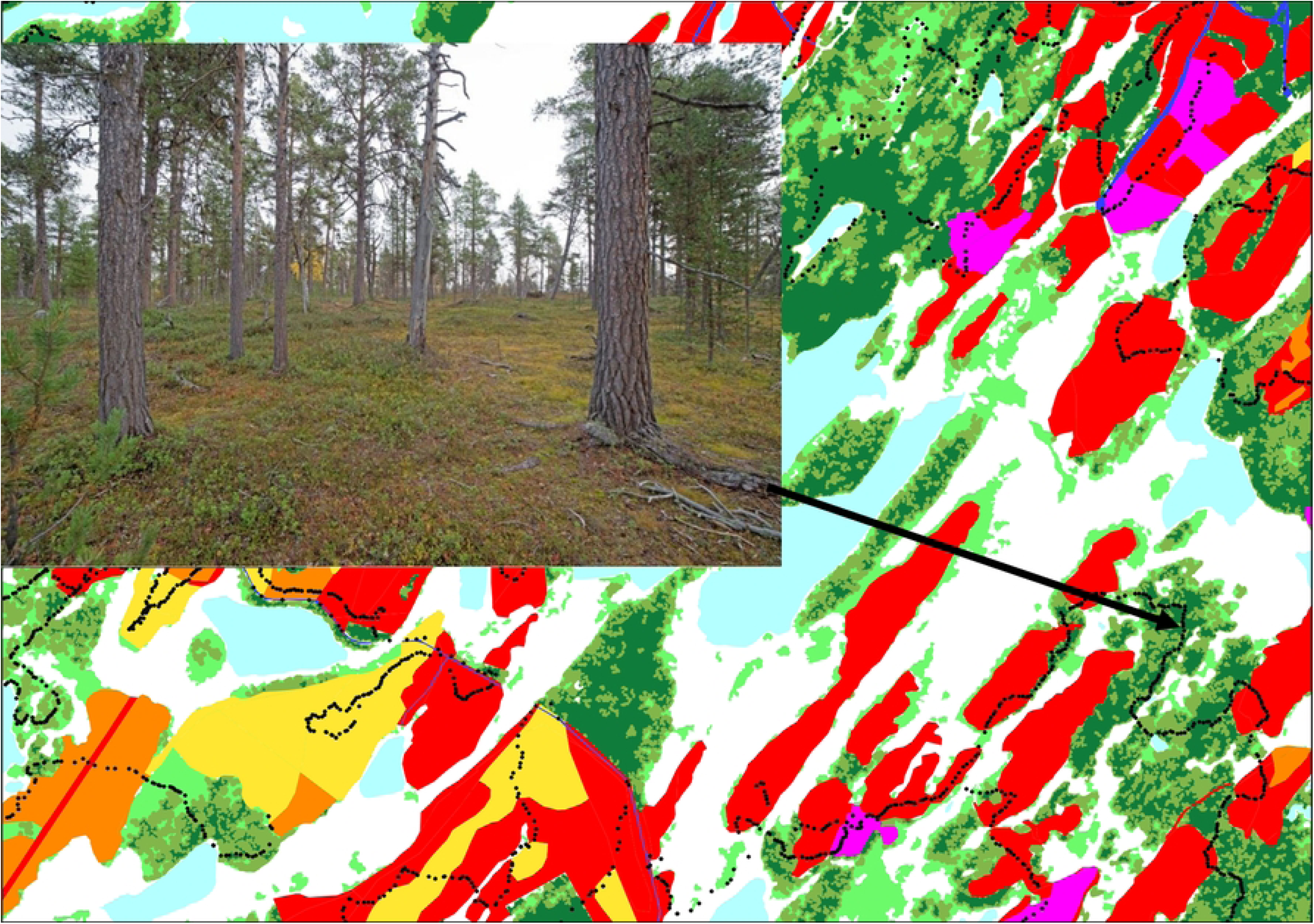
Calibrating stand types using 52 000 GPS-referred photos.

### Implementing the lichen simulation with model variations

The lichen equations biomass, growth and usable growth equations were implemented in a python code reading the shape-file database dbf-file. The forest change was analyzed in 5-year periods as the reindeer amount of previous 5-year period correspond well for the lichen biomass equations (1) and (2). All lands of Mudusjävri co-operative were divided into the required pasture classes of the static lichen model. Table 3 shows the pasture classes with lichen and the mapped forests development classes defined into these eleven pasture classes:

1. Dry logging areas and sapling stands
2. Semi-dry logging areas and sapling stands
3. Dry young pine forest (31 - 80 years)
4. Semi-dry young pine forest (31 - 80 years)
5. Dry mature pine forest (81 - 140 years)
6. Semi-dry mature pine forest (81 - 140 years)
7. Dry old pine forest (> 140 years)
8. Semi-dry old pine forest (> 140 years)
9. Dry mountain birch forests
10. Semi-dry mountain birch forests
11. Mountain heaths

For the digitized forest areas, the trophic level was achieved from the Luke and Syke pasture class raster data. The development class was achieved from our mappings by the categories of table 3. The amount of arboreal lichen forests was calculated by summing the areas of mature and old pine forests.

**Table 3.** Forest Pasture classification.

| Pasture class | Corresponding mapped forest development class |
| --- | --- |
| Logging areas and sapling stands | - Logged stands<br>- Logging stands with saplings when less than 30 years from loggings<br>- Old logging with young trees in clusters when less than 30 years from loggings |
| Young pine forest | - Natural low canopy forests<br>- Thinned young forest<br>- Logging stands with saplings when more than 30 years from loggings |
| Mature pine forest | - Old logging with young trees in clusters when over 30 years after loggings |
| Old pine forest | - Natural, dense, high canopy<br>- Natural, mixed size trees<br>- Thinned forest stand<br>- Cautious selective logging |

Pasture classification of mapped forest development classes.

As arboreal lichen is an important winter fodder also the ground lichen models defined by equations (1) – (4), the annual arboreal lichen production was added to these models. According to previous publications the amount of arboreal lichen in Inari scots pine forests in typical sub dry sites as in Mudusjävri co-operative varies between 40 to 400 kg/ha depending much on the stand volume [24]. In typical old pine forests with volumes over 100 m3/ha the mean amount of arboreal lichen is 23 kg/ha and 120 kg/ha in dry and sub-dry sites, respectively. From this only 1 % is below 2 m height reachable for the reindeer but litterfall during winter gives the most reachable arboreal lichen being 0.1 - 22 kg/ha/year [31]. Thus, it can be assumed that old pine forest produces in average about 11 kg/ha/year arboreal lichen to the reindeer to feed in sub-dry old and mature forests. Thus, it can be estimated that the amount of reachable arboreal lichen is about 9 % of the biomass. In our models a 50 % loss in foraging was assumed resulting in a usable annual arboreal amount approximately 4.5 % of the total lichen biomass on the trees.

#### Model 1 – the baseline model

The baseline model used in ground lichen simulation was the basic model reported in [15] and [16] as such. The reindeer amount was kept constant with the average amount of years 2000 – 2010.

#### Model 2

The baseline model was used with the real reindeer amounts of 1950 to 2020.

#### Model 3

The baseline model pasture class parameter was updated with the values of Table 1 of the latest pasture inventory. The reindeer amount was kept constant as in model 1.

#### Model 4

Model 3 was varied by simulating with the real reindeer amounts of 1950 to 2020.

#### Model 5

Traditional pasture class knowledge of reindeer herders was taken into account in the trophic level of forest stands. An assumption was made that the good lichen areas before industrial loggings were dry sites and turned into semi-dry stands with denser vegetation as forest has grown. A gradual change from dry to semi-dry in 30 years after loggings was implemented. Model 5 used a constant reindeer amount of the average of years 2000 – 2010.

#### Model 6

Model 5 was varied by simulating with real reindeer amounts of 1950 to 2020.

## Results

We start the presentation of the results with a map for year 2020 (Fig 6), which is based on the categories of different forest stands according to their age and structure. Different shades of the colour green represent forests in natural state, and the map shows that only a minority of the forests remain intact by forestry today. A lot of the natural low canopy forests in the border area of summer and winter grazing areas are tundra birch forests and not suitable for economic forestry. Approximately 60 % of areas suitable for forestry has been logged during the period of industrial loggings that started in the mid-1950s. 25 200 ha or 45 % of all forests have been logged. Before Mid-1950s all forests were practically in natural state as only some insignificant amounts of selective loggings had been done for local huts and buildings.

**Fig 6.**
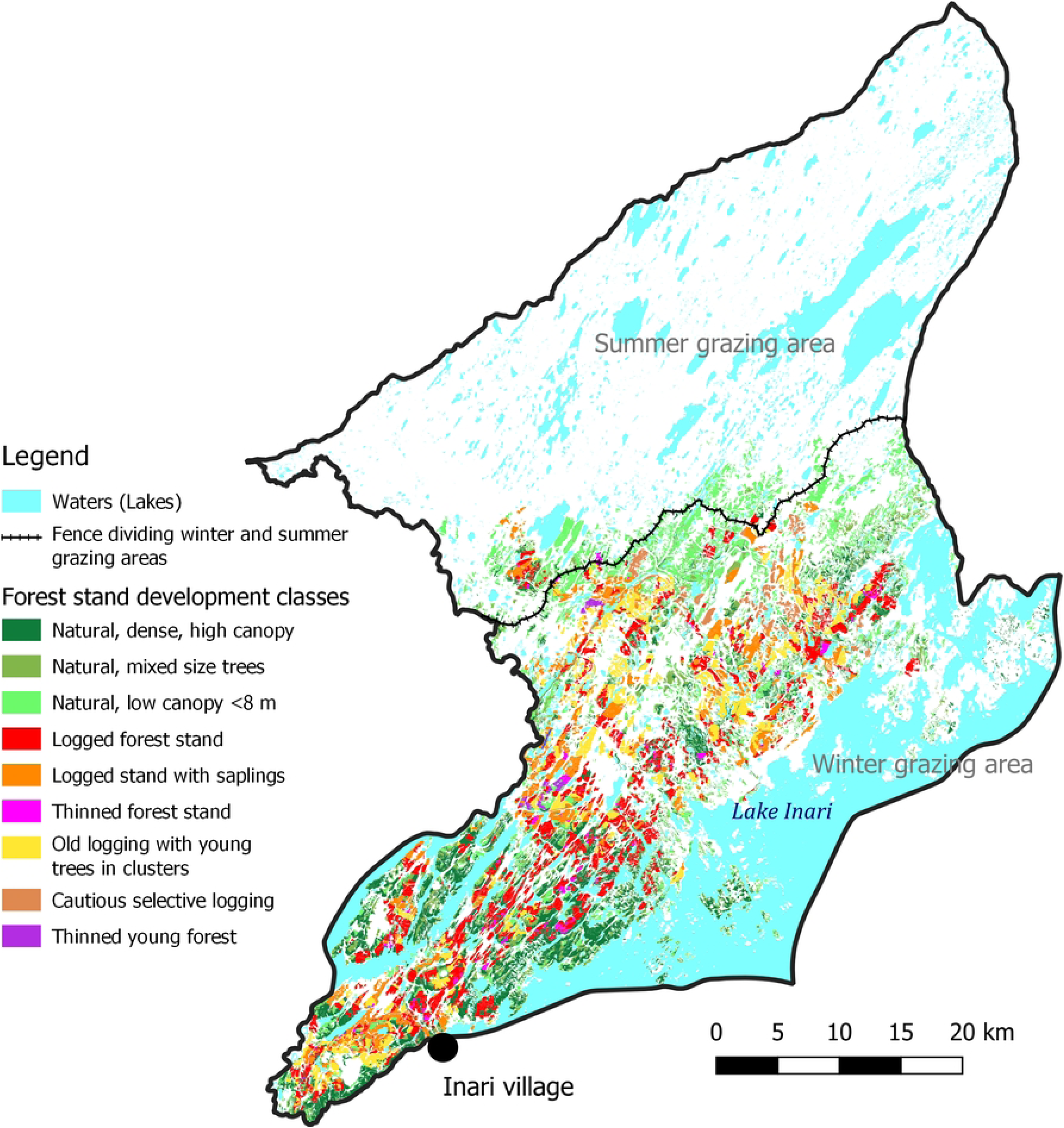
Forest stands on productive forest land of Mudusjävri co-operative’s winter pasture area.

The pasture class lichen biomasses for year 2020 are shown in Fig 7. for the winter grazing areas. Model 5 was used in this example. The summer grazing area is similar but with values being approximately 30 % for ground lichen. The differences for year 2020 vary very little between the different model variations. Arboreal biomasses were taken as such from reference [24]. The model gives an average ground lichen biomass of 222 kg/ha and 63 kg/ha for the winter and summer grazing areas, respectively. According to the latest inventory for years 2016 – 2018 [34], the respective values are approximately 230 kg/ha and 80 kg/ha for winter and summer grazing areas, respectively. The calculated average biomasses seem to match the inventory values rather well.

**Fig 7.**
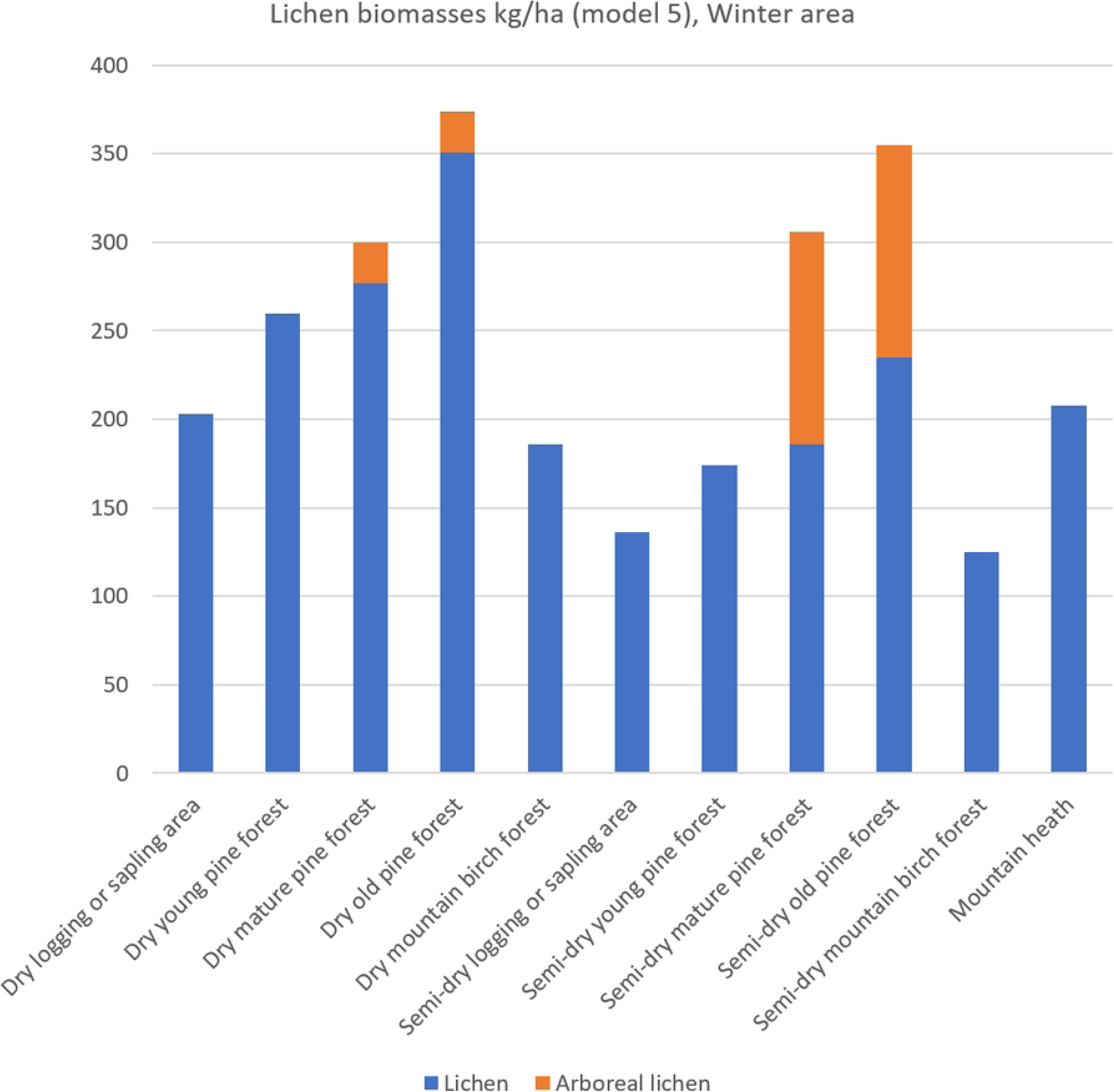
Ground and arboreal lichen biomasses in Mudusjävri winter grazing area of year 2020 using model 5.

The corresponding yearly lichen growth in year 2020 is shown in Fig 8. This is, however, not the amount of lichen the reindeer can use or forage. The effective growth usable for reindeer in a year is shown in Fig 9. The reindeer’s preference of mature and old pine forests is quite understandable when the loggings or young forests have less than one third to half of the usable lichen compared to mature and old forests. Thus, figures suggest also severe impacts of forestry. Loggings in old forests reduce the usable lichen production by 72 – 79 %.

**Fig 8.**
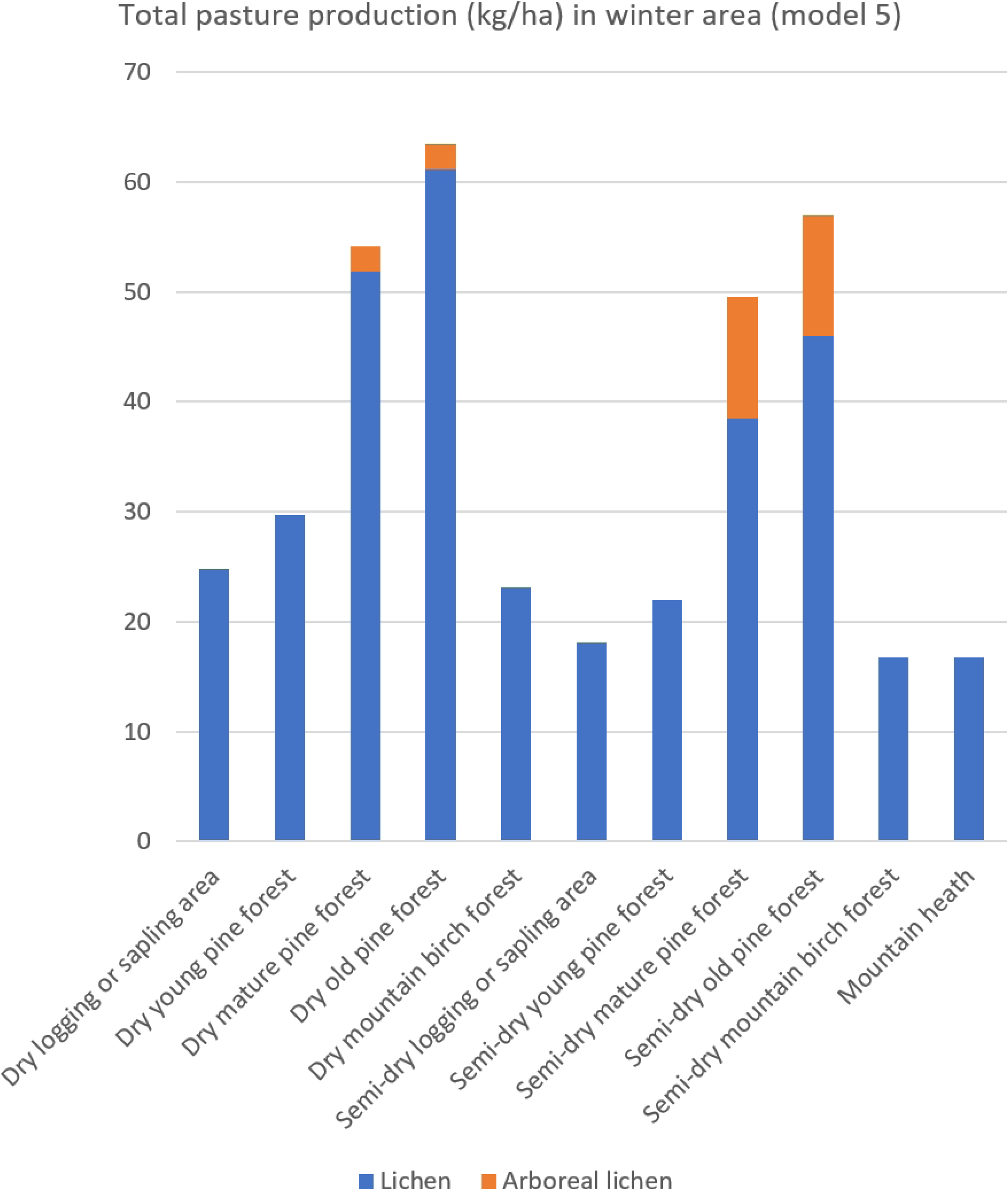
Total lichen growth in Mudusjävri winter grazing area in 2020 according to model 5.

**Fig 9.**
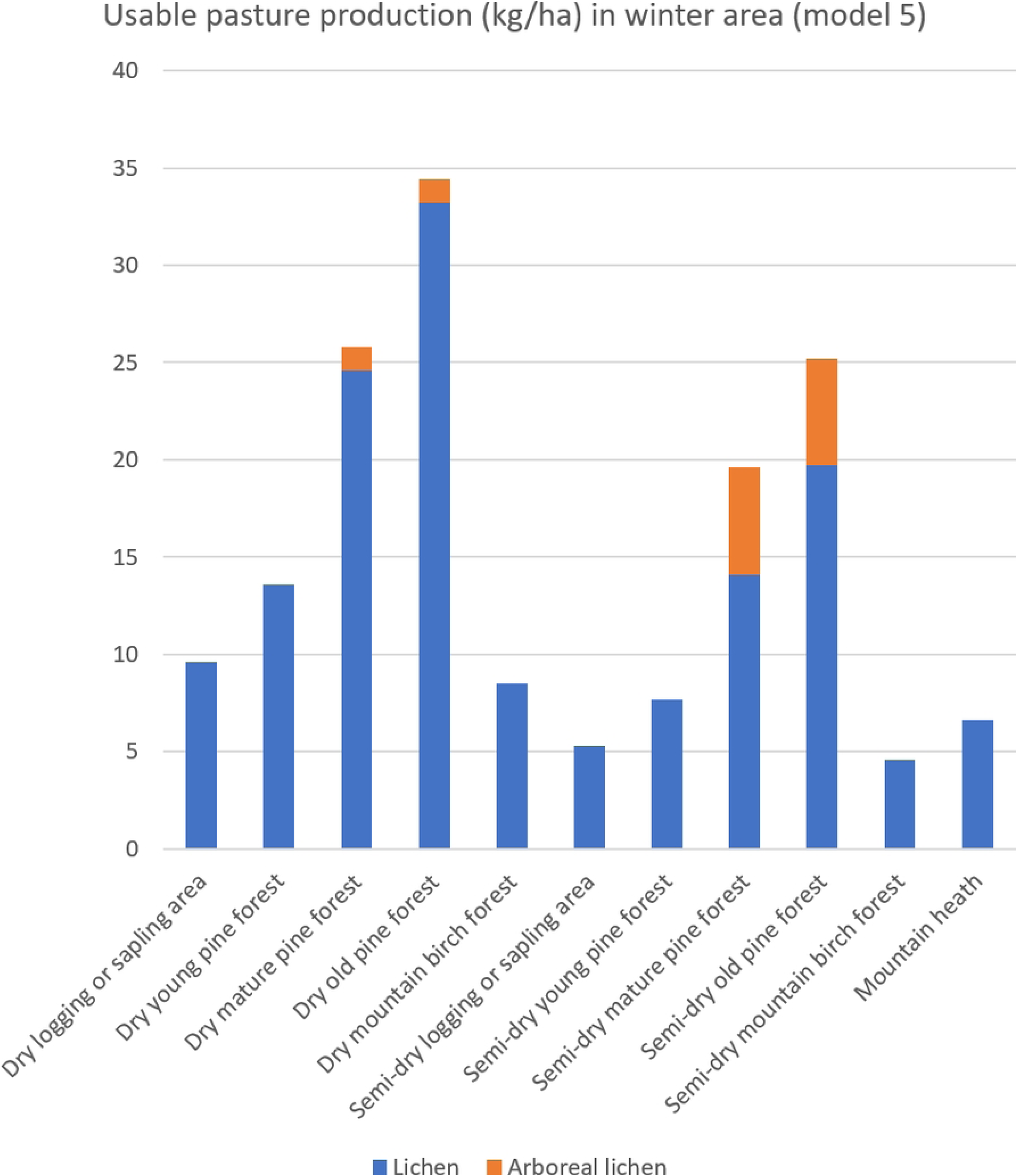
Usable pasture production in the winter grazing area in Mudusjävri co-operative according to model 5.

To have a historical reference for impact assessment, the lichen amounts of year 1950 were chosen as the baseline. At that time there had been no large-scale industrial loggings. The model variations 1 to 6 were simulated from 1950 to 2020 with the mapped forest structure change. The resulting usable lichen amounts are shown in Fig 10. Depending on the model the usable ground lichen amount was around 1900 – 2200 tons/year in 1950 whereas forest change has decreased the amounts to 1100 – 1200 tons/year in 2020. Absolute decrease in usable lichen growth is 680 - 1020 tons/year depending on the model. The relative decrease of usable ground lichen growth due to forest change is 36 – 47 % depending on the model. The radical increase in usable lichen production between 1970 – 1980 by the model is due to leap year in reindeer herding in the beginning of 1970s. Tough winter and snow conditions caused a huge drop in reindeer amounts.

**Fig 10.**
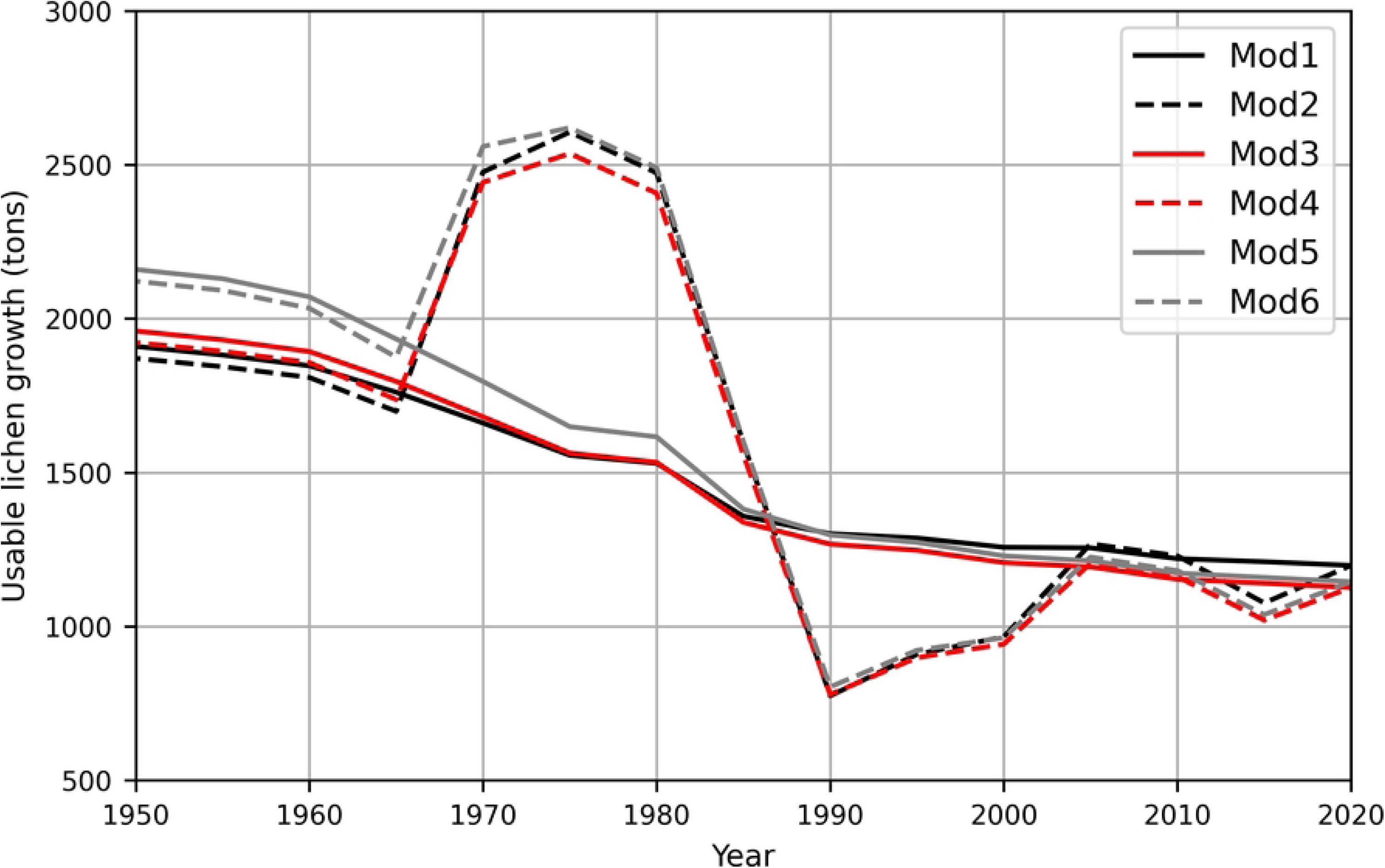
Estimated usable ground lichen growths in Mudusjävri region during 1950 - 2020.

Although the loggings were in quite a large level in the 1990s the usable lichen growth decreases slowly as the large loggings of 1960s starts to change into more productive pasture classes.

The cumulative economic costs for Mudusjävri reindeer herding can be estimated with different approaches. We chose to calculate the respective feeding cost if the usable lichen growth losses of Fig 10. are compensated with artificial feeding. The current feeding costs are approximately 0.5 €/kg [48]. The resulting plot of feeding costs are shown in Fig 11. The feeding costs increase constantly due to cumulative logging impacts from 1950 onward for the models with constant reindeer amounts. For the models with real reindeer amounts the costs are negative due to the small reindeer amounts in the 1970s. Depending on the model used, the yearly additional feeding costs in 2020 are approximately from 370 000 to 530 000 €/year. During period 2010 – 2020 the actual costs of the additional feeding has been approximately 140 000 €/year according to the accounting of Mudusjävri co-operative. According to the reindeer herders these costs would not exist without pasture losses and the co-operative region would support a larger number of reindeer and would result into larger incomes for the reindeer herders. According to our simulations, these claims seem credible.

**Fig 11.**
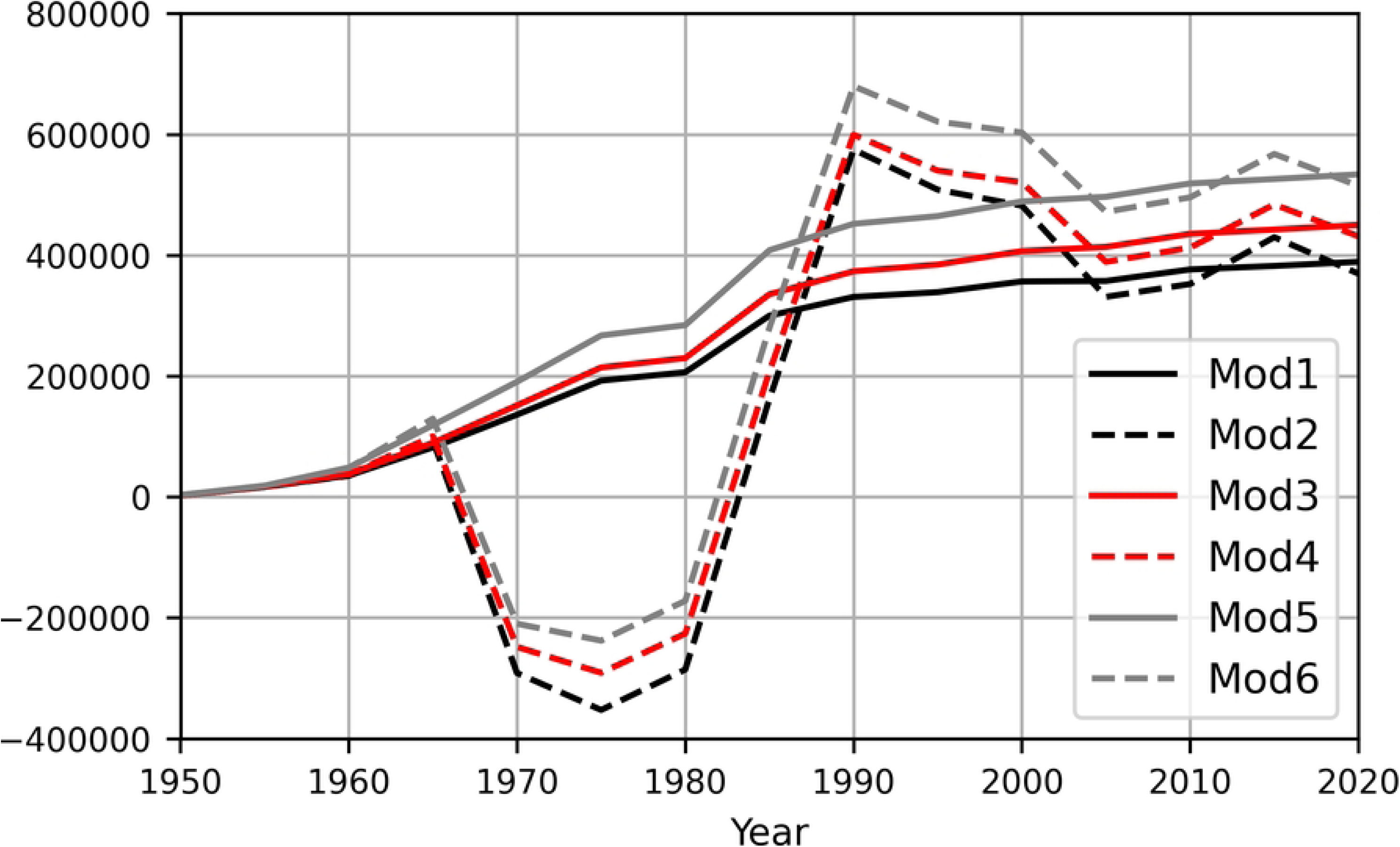
Estimated economic losses of Mudusjävri reindeer herding co-operative due to forestry.

Due to the infrastructure disturbance effect of the lichen biomass model (or eq. (2)) the forest road effect can be estimated separately. The forestry roads increased rather steadily from 1950 to 1990 from 0 to 200 km. According to model variation 5 the forest road infrastructure causes a cost of 50 000 €/year. This is approximately about 250 €/km/year. Similarly, infrastructure disturbance area analysis of cabins shows a cost effect of 20 000 €/year.

The amount of arboreal lichen loss in year 2020 correspond to loss of 31 000 euro/year in models 1 to 4. The forests would feed the reindeer annually with 61 tons more arboreal lichen with the forest structure of the 1950s. In models 5 ja 6, where the dry to sub-dry conversion was assumed in the forests defined by reindeer herders, the arboreal lichen loss would be 46 tons/year corresponding to an artificial feeding cost of 23 000 euro/year. These values are rather small compared to the ground lichen losses. However, arboreal lichen can be crucial in spring times with much hard snow when it is the only nutrition, they are able to reach and ground lichen is hard to dig.

**Fig 12.**
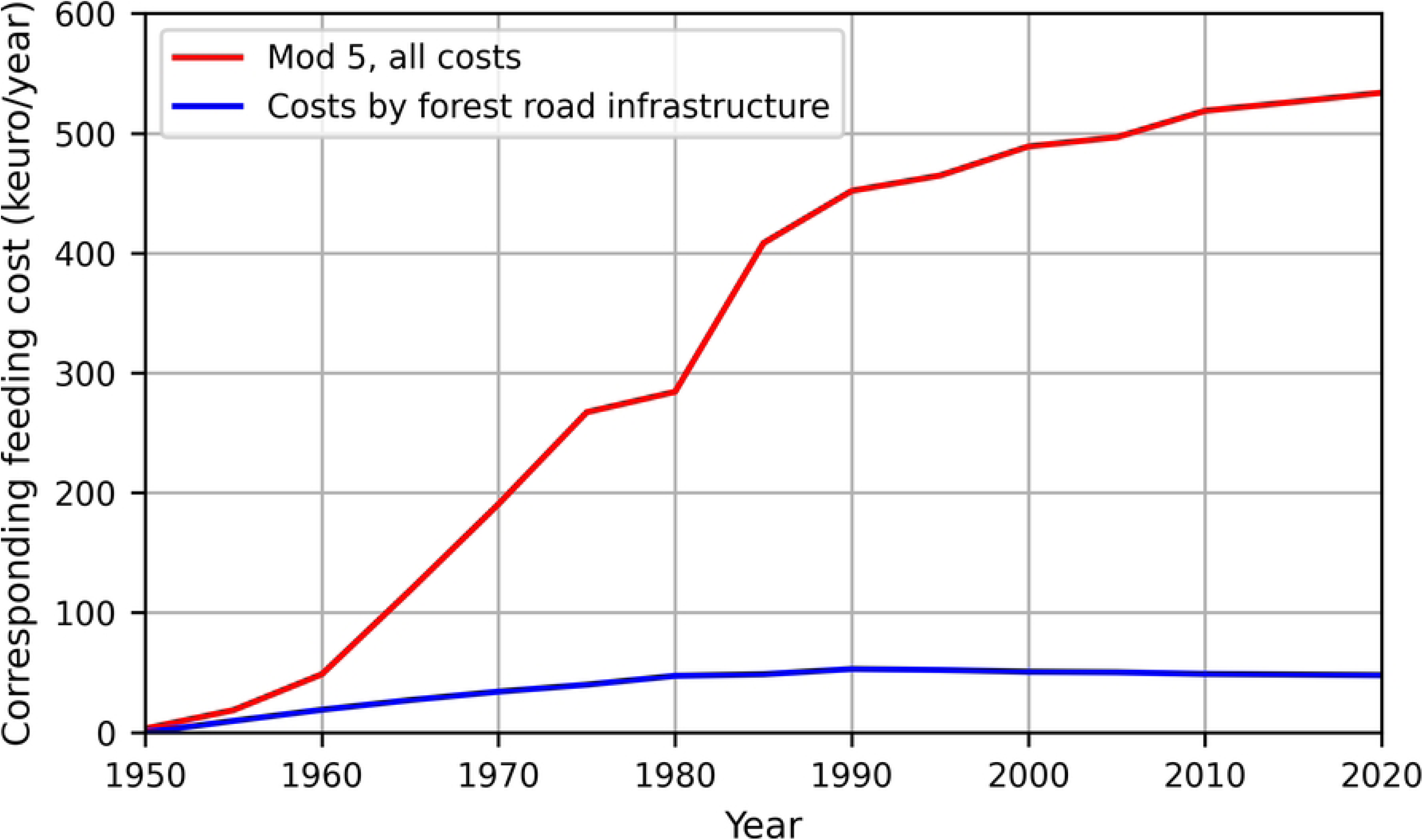
Estimated feeding costs of road infrastructure according to model 5.

The feed requirement of reindeer is approximately 2 kg ground lichen per reindeer per day [49]. The winter herd size in Mudusjävri co-operative is at maximum 5 200 reindeer, which means that the simulated 740 - 1 070-ton loss of feed corresponds to a 70 - 100-day need of the entire winter herd. Thus, 2 - 3 months loss in nutrition is caused for the Mudusjävri winter herd by the changes in forest structure caused by forestry dependent on the model used.

Another approach to assess economic impacts on reindeer herding could be to estimate the sustainable herd size winter pastures enable and calculate the respective meat production loss. Mudusjävri accounting shows that in typical years during the last decade gives a 600 000 € revenue of slaughtering of reindeer for the winter herd of 5200 reindeer (approx. 115 €/reindeer/year). Ground lichen is the most important winter fodder for reindeer and practically define the sustainable number of reindeer. If the current winter herd of 5200 is considered sustainable determined from the pasture inventory [34] it can be estimated that with the untouched forest structure of 1950s the reindeer number could be 8100. This is calculated from our more cautious model 1 giving a 36 % reduction in usable lichen growth. Thus, the meat production revenue reduction for the lost 2900 reindeer is about 340 000 €/year. This is rather comparable to the model 1 feeding cost estimation of 390 000 €/year.

## Discussion

Our lichen growth simulation gives the minimum level of harm caused by forestry impacts and only on the amount of ground and arboreal lichen production. However, there are many other effects of forestry not taken into account in these calculations. According to our interviews of Mudusjävri reindeer herders some of the most severe impacts are:

1. Reindeer prefer old forest over young in winter time. Large amount of young forests makes the controlling of the herd difficult.
2. Logged stands or sparse sapling stands tend to have harder snow due to different wind conditions. Digging of ground lichen is more difficult for the reindeer
3. Scattering of pastures makes unconnected smaller old forest stands less valuable as winter pastures. Reindeer herds tend to graze in connected old forests during winter.
4. Scattered pastures make reindeer wander more and some places require uniform unshattered forest areas to stop reindeer from migrating to other co-operatives.
5. Logging residues drive reindeer away even though there still would be ground lichen left, because they cannot access the lichen that is covered.
6. Valuable pastures may not be possible to use due to disturbance by e.g., large predators or by human activity such as tourism.

These are partly the same as previously mentioned in Introduction chapter. All these effects result into more herding work for the reindeer herders increasing their costs. In other words, the usability of the different stands and grazing areas have not been taken into account in this study.

Many other choices were carefully made not to overestimate forestry impacts. The mapping accuracy is a source of error and many forest categories were estimated to be better than they most likely are due to the lack of more accurate information. Thinned stands were assumed to be as good lichen areas as natural forests which is very likely not the case for the time there are logging residues. According to Mudusjävri reindeer herders the logging residues make reindeer avoid the area during winter time for a period varying from 4 - 15 years depending on stand type and forestry activities.

The annual revenue of optimal pasture ha is according to [50] 30 €/ha. This number is determined from the meat production with optimal pastures. For comparison, it is interesting to note that the annual revenue calculated from lichen growth production is in the same magnitude of order. In Mudusjävri old pine forests stands the total available annual lichen production is approximately 25 to 34 kg/ha which correspond to an annual feeding cost of 13 to 17 €/ha. This is rather well in line with the order of magnitude of revenue of optimal pasture conditions of [50].

Although, there are uncertainty in the method and lichen amounts, we conclude that we have credibly assessed that forestry has caused significant harm to Sámi culture by deteriorating pasture conditions of Mudusjävri co-operative reindeer herding livelihood. This outcome bases solely on lichen amount assessment and by omitting all the other detrimental forestry impacts on reindeer herding, such as usability decrease by shattering. Combining the forestry impacts on reindeer herding – combined with its low economic feasibility in Inari region [51] - the justification of industrial loggings in the recent levels seem highly questionable.

## Conclusions

In this paper we have presented the methodology for a map-based impact assessment method that was developed to assess the impacts of forestry on reindeer grazing areas, and the results of such a mapping in the pasture areas of Mudusjävri reindeer herding co-operative. We argue that this type of impact assessment is both a relevant and necessary way to measure the significance of the harm caused by forestry to Sámi culture and livelihoods, as protected by domestic and international law. Further, based on our analysis, we conclude that the harm caused to Mudusjävri co-operative must be deemed as significant. Our model suggests that logging a stand with old-growth pine forest results in a 70 - 80 % reduction in usable ground lichen growth of that stand for a period of at least 30 years. According to lichen simulation the change in forest structure from 1950 to 2020 has, based on the lichen model variation used, reduced the usable lichen growth in Mudusjävri reindeer herding co-operative by 36 - 47 %, corresponding to a decrease in lichen growth of 680 - 1020 tons/year and, consequently, an economic loss of 370 000 – 530 000 €/year for reindeer husbandry in the co-operative if current artificial feeding costs are used and the loss was fully compensated for by feeding. Having now established the significance of the impacts, the next step in our work is to formulate a forest management plan for the Mudusjävri co-operative, jointly with the reindeer herders. The goal of the plan is to describe forest management methods to specified regions that would improve the quality the winter pastures by through a combination of restrictions to further loggings and active restoration measures.

## Acknowledgments

This work is part of “What form can an atonement take?” -project (*Miltä sopu näyttää*). None of this work would have been possible without the reindeer herders of Mudusjävri reindeer herding co-operative, and their families. We are indebted to their indigenous knowledge and their commitment to our collaborative work. Our warmest thanks go also to Juha Länsman for his extraordinary effort in producing the photo data set.

## References

1. IPBES. Summary for Policymakers of the Global Assessment Report on Biodiversity and Ecosystem Services of the Intergovernmental Science-Policy Platform on Biodiversity and Ecosystem Services; Díaz, S., Settele, J., Brondizio, E.S., Ngo, H.T., Guèze, M., Agard, J., Arneth, A., Balvanera, P., Brauman, K.A., Butchart, S.H.M., Eds.; IPBES Secretariat: Bonn, Germany, 2019.

2. Steffen W, Broadgate W, Deutsch L, Gaffney O, Ludwig C. The trajectory of the Anthropocene: The Great Acceleration. The Anthropocene Review. 2015;2(1):81–98. doi:10.1177/2053019614564785

3. Åhrén M. Indigenous peoples’ status in the international legal system. Oxford: University Press; 2016: 202ff.

4. Anaya J, Report of the Special Rapporteur on the situation of human rights and fundamental freedoms of indigenous people, United Nations Human Rights Council (A/HRC/12/34, 2009).

5. Larsen RK. Impact assessment and indigenous self-determination: a scalar framework of participation options, Impact Assessment and Project Appraisal. 2017. doi: 10.1080/14615517.2017.1390874.

6. Larsen RK, Raitio K, Stinnerbom M, Wik-Karlsson J. Sami-state collaboration in the governance of cumulative effects assessment: A critical action research approach. Environ. Impact Assess. Rev. 2017; 64: 67–76.

7. Weber M, Krogman N, Antoniuk T. Cumulative Effects Assessment: Linking Social, Ecological and Governance Dimensions. Ecology and Society. 2012; 17; 2: Available from: https://www.jstor.org/stable/26269026

8. Kivinen S. Many a little makes a mickle: Cumulative land cover changes and traditional land use in the Kyrö reindeer herding district, northern Finland. Appl. Geogr. 2015; 63: 204–211.

9. Österlin C, Raitio K. Fragmented Landscapes and Planscapes—The Double Pressure of Increasing Natural Resource Exploitation on Indigenous Sámi Lands in Northern Sweden. Resources 2020. 2020; 9: 104. doi: 10.3390/resources9090104

10. Brännlund I, Axelsson P. Reindeer management during the colonization of Sami lands: A long-term perspective of vulnerability and adaptation strategies. Global Environmental Change 21. 2011; 1095–1105. doi: 10.1016/j.gloenvcha.2011.03.005.

11. Nicholas JC, Tyler IHB, Førland EJ, Nellemann C. The Shrinking Resource Base of Pastoralism: Saami Reindeer Husbandry in a Climate of Change. Frontiers in Sustainable Food Systems. 2021; 10. Doi: 10.3389/fsufs.2020.585685

12. Larsen RK, Österlin C, Guia L. Do voluntary corporate actions improve cumulative effects assessment? Mining companies’ performance on Sami lands. The Extractive Industries and Society. Elsevier. Forthcoming. doi: 10.1016/j.exis.2018.04.003.

13. Saarikoski H, Raitio K. Science and politics in old-growth forest conflict in Upper Lapland. Nature and Culture 8. 2013; 1:53–73.

14. Raitio K. Seized and missed opportunities in responding to conflicts: constructivity and destructivity in forest conflicts management in Finland and British Columbia, Canada. In: Peterson TR, Ljunggren Bergeå H, Feldpausch-Parker AM, Raitio K. (Eds): Environmental Communication and Community. Constructive and destructive dynamics of social transformation. Routledge Earthscan. 2016; 229–249.

15. Kumpula J, Kurkilahti M, Helle T, Colpaert A, Both reindeer management and several other land use factors explain the reduction in ground lichens (Cladonia spp.) in pastures grazed by semi-domesticated reindeer in Finland. Reg Environ Change. 2014; 14: 541 – 559. doi: 10.1007/s10113-013-0508-5

16. Pekkarinen AJ, Kumpula J, Tahvonen O. Reindeer management and winter pastures in the presence of supplementary feeding and government subsidies. Ecological Modelling. 2015; 312: 256 – 271. doi: 10.1016/j.ecolmodel.2015.05.030

17. Kojola I, Helle T, Niskanen M, Aikio P. Effects of lichen biomass on winter diet, body mass and reproduction of semi-domesticated reindeer Rangifer t. tarandus in Finland, Wildlife Biology. 1995; 1: 33 – 38

18. Helle T, Tarvainen L. Determination of winter digging period of semi-domesticated reindeer in relation to snow conditions and food resources. - Reprints of Kevo Subarctic Research Station. 1984; 19: 49 – 56

19. Jokinen M. Heated and frozen forest conflicts: Cultural sustainability and forest management in arctic Finland. IUFRO World Series. Vienna. 2014; 32. 561p.

20. Vihervaara P, Kumpula T, Tanskanen A, Burkhard B. Ecosystem services–A tool for sustainable management of human–environment systems. Case study Finnish Forest Lapland. Ecological Complexity. 2010; 7: 410–420.

21. Rytkönen AM, Saarikoski H, Kumpula J, Hyppönen M, Hallikainen V. Metsätalouden ja poronhoidon väliset suhteet Ylä-Lapissa– synteesi tutkimustiedosta. Metlan työraportteja / Working Papers of the Finnish Forest Research Institute 6/2013. Available from: https://jukuri.luke.fi/bitstream/handle/10024/504425/rkts2013_6.pdf. Finnish

22. Esseen PA, Renhom KE, Pettersson R. Epiphytic lichen biomass in managed and old-growth boreal forests: Effect of branch quality. Ecological Applications. 1996; 6: 228–238

23. Anttonen M, Kumpula J, Colpaert A. Range selection by semi-domesticated reindeer (Rangifer tarandus tarandus) in relation to infrastructure and human activity in boreal forest environment, northern Finland. Arctic 64. 2011; 1: 1–14

24. Jaakkola LM, Helle TP, Soppela J, Kuitunen MT, Yrjönen MJ. Effects of forest characteristics on the abundance of alectorioid lichens in northern Finland. Canadian Journal of Forest Research. 2006; 36: 2955–296. doi:10.1139/X06-178

25. Helle T, Aspi J, Kilpela SS. The effects of stand characteristics on reindeer lichens and range use by semidomesticated reindeer. Rangifer Special Issue 3; 1990; 3: 107–114

26. Larsen R, Raitio K, Sandström P, Skarin A, Stinnerbom M, Wik-Karlsson J et al. Kumulativa effekter av exploateringar på renskötseln. Naturvårdsverket; 2016. Swedish

27. Heinämäki L. Saamelaiskulttuurin heikentämiskielto ja viranomaisten aktiivinen velvoite turvata perinteisten elinkeinojen harjoittamisen ja kehittämisen edellytykset. Lakimies. 2021; 1: 3 – 35. Finnish

28. Heinämäki L. Opas saamelaisia koskevien oikeusnormien tulkintaan ja soveltamiseen ympäristöön ja maankäyttöön liittyvissä kysymyksissä. Saamelaiskäräjät / Saami parliament of Finland; 2021. Finnish

29. Landauer M, Rasmus S, Forbes BC. 2021. What drives reindeer management in Finland towards social and ecological tipping points? Regional Environmental Change. 2021; doi: 10.1007/s10113-021-01757-3

30. Kumpula J, Pekkarinen AJ, Tahvonen O, Rasmus S. Poronhoidon tuottavuus ja ekonomia erilaisissa laidun-ja ympäristöolosuhteissa. Yhteenveto tutkimushankkeesta. Luonnonvara-ja biotalouden tutkimus 68/2015. Luonnonvarakeskus. Helsinki. 2015; Available from: http://urn.fi/URN:ISBN:978-952-326-144-0. Finnish

31. Mattila E. Survey of reindeer winter ranges as a part of the Finnish National Forest Inventory in 1976–1978. Communicationes Instituti Forestalis Fenniae 99. 1981; 6: 74

32. Mattila E. Porojen talvilaitumien kunto Ylä-Lapin paliskunnissa vuonna 2004. Metlan työraportteja / Working Papers of the Finnish Forest Research Institute 28. 2006; Available from: http://www.metla.fi/julkaisut/workingpapers/2006/mwp028.htm. Finnish

33. Mattila E. Ylä-Lapin talvilaidunarvioinnin tuloksia. Uusimmat arviot vuodelta 2012 ja vastaavia tuloksia vuodelta 2004. Metlan työraportteja / Working Papers of the Finnish Forest Research Institute 282. 2014; Available from: http://www.metla.fi/julkaisut/workingpapers/2014/mwp282.htm. Finnish

34. Kumpula J, Siitari J, Siitari S, Kurkilahti M, Heikkinen J, Oinonen K. Poronhoitoalueen talvilaitumet vuosien 2016–2018 laiduninventoinnissa: Talvilaidunten tilan muutokset ja muutosten syyt. Luonnonvara-ja biotalouden tutkimus 29/2019. Luonnonvarakeskus. Helsinki. 2019; Available from: https://jukuri.luke.fi/bitstream/handle/10024/544124/luke-luobio_33_2019.pdf. Finnish

35. Kumpula J, Colpaert A, Anttonen M. Does forest harvesting and linear infrastructure change the usability value of pastureland for semi-domesticated reindeer (Rangifer tarandus tarandus)? Ann. Zool. Fennici. Helsinki. 2007; 44: 161 – 178

36. Kivinen S, Moen J, Berg A, Eriksson Å. Effects of Modern Forest Management on Winter Grazing Resources for Reindeer in Sweden. Ambio. 2010; 39(4): 269–278, doi: 10.1007/s13280-010-0044-1

37. The public GIS-data download service of Syke, Finnish Environment Institute: https://www.syke.fi/fi-FI/Avoin_tieto/Paikkatietoaineistot/Ladattavat_paikkatietoaineistot

38. The public GIS-data download service of National Land Survey of Finland. Available from: https://tiedostopalvelu.maanmittauslaitos.fi/tp/kartta?lang=en

39. The public GIS-data download service of Metsäkeskus Forest Centre. Available from: https://www.metsaan.fi/paikkatietoaineistot

40. Vilhomaa J, Laaksonen H. Valtakunnallinen laserkeilaus – Testityöstä tuotantoon. The Photogrammetric Journal of Finland. 2011; 22. No. 3: 82 – 91

41. QGIS www-information and download source of the open-source GIS-software. Available from: https://qgis.org/

42. GRASS GIS www-information and download source of the open-source GIS-software. Available from: https://grass.osgeo.org/

43. Landsat imagery download service. Available from: https://glovis.usgs.gov/

44. Sentinel satellite imagery download service. Available from: https://scihub.copernicus.eu/

45. Sandström O, Vaara I, Heikkuri P, Jokinen M, Kokkoniemi T, Liimatainen J. et al. Ylä-Lapin luonnonvarasuunnitelma. Metsähallituksen metsätalouden julkaisuja 38. ISBN 952-446-253-2. 2000. Finnish

46. Lawrence R, Larsen RK. Fighting to be herd, Impacts of the proposed Boliden copper mine in Laver, Älvsbyn, Sweden for the Semisjaur Njarg Sami reindeer herding community. Stockholm Environment Institute report. 2019; Available from: https://www.sei.org/wp-content/uploads/2019/04/sei-report-fighting-to-be-herd-300419.pdf

47. Larsen RK, Boström M, Muonio Reindeer Herding District, Vilhelmina Södra Reindeer Herding District, Voernese Reindeer Herding District, Wik-Karlsson. The impacts of mining on Sámi lands: A knowledge synthesis from three reindeer herding districts. The Extractive Industries and Society. 2022 March; 9: 101051. Available from: https://www.sciencedirect.com/science/article/pii/S2214790X22000090

48. Kumpula J, Siitari S. Kestävä biotalous porolaitumilla -hankkeen osaraportit, johtopäätökset ja toimenpide-ehdotukset, Luonnonvara-ja biotalouden tutkimus 29/2020, Luke. 2020. Available from: https://jukuri.luke.fi/bitstream/handle/10024/545810/luke_luobio_29_2020.pdf

49. Nieminen M, Pokka AS, Heiskari U. Artificial feeding and nutritional status of semi-domesticated reindeer during winter. Rangifer. 1987; 7 (2): 51 – 58. doi: 10.7557/2.7.2.717

50. Tahvonen O, Kumpula J, Pekkarinen AJ. Optimal harvesting of an age-structured, two-sex herbivore-plant system. Ecological Modelling. 2014; 272: 348 – 361

51. Parkatti VP, Tahvonen O. Economics of Multifunctional Forestry in the Sámi People Homeland Region. FEEM Working Paper No. 25. 2020; Available from: https://ssrn.com/abstract=3749963

